# OpenAnnotate: a web server to annotate the chromatin accessibility of genomic regions

**DOI:** 10.1101/596627

**Authors:** Shengquan Chen, Qiao Liu, Xuejian Cui, Zhanying Feng, Chunquan Li, Xiaowo Wang, Xuegong Zhang, Yong Wang, Rui Jiang

## Abstract

Chromatin accessibility, as a powerful marker of active DNA regulatory elements, provides valuable information for understanding regulatory mechanisms. The revolution in high-throughput methods has accumulated massive chromatin accessibility profiles in public repositories. Nevertheless, utilization of these data is hampered by cumbersome collection, time-consuming processing, and manual chromatin accessibility (openness) annotation of genomic regions. To fill this gap, we developed OpenAnnotate (http://health.tsinghua.edu.cn/openannotate/) as the first web server for efficiently annotating openness of massive genomic regions across various biosample types, tissues, and biological systems. In addition to the annotation resource from 2729 comprehensive profiles of 614 biosample types of human and mouse, OpenAnnotate provides user-friendly functionalities, ultra-efficient calculation, real-time browsing, intuitive visualization, and elaborate application notebooks. We show its unique advantages compared to existing databases and toolkits by effectively revealing cell type-specificity, identifying regulatory elements and 3D chromatin contacts, deciphering gene functional relationships, inferring functions of transcription factors, and unprecedentedly promoting single-cell data analyses. We anticipate OpenAnnotate will provide a promising avenue for researchers to construct a more holistic perspective to understand regulatory mechanisms.

## INTRODUCTION

Chromatin accessibility is a measurement of the ability of nuclear macromolecules to physically contact DNA and is a critical determinant of understanding regulatory mechanisms (1). For example, accessible genomic regions are regarded as the primary positions of regulatory elements (2), and provide a great opportunity to study transcription factor binding sites, DNA methylation sites, histone modifications, gene regulation, and regulatory networks (3,4). Besides, changes in chromatin accessibility have been implicated with different perspectives of human health as a result of the alterations of nucleosome positioning affected by mutations in chromatin remodelers (5–7). A number of high-throughput technologies have been developed to profile chromatin accessibility, such as ATAC-seq (8), DNase-seq (9), FAIRE-seq (10), and MNase-seq (11). DNase-seq was widely applied in consortiums such as ENCODE (12,13), while ATAC-seq, as a more recent technique, can provide higher accuracy and sensitivity. Both DNase-seq and ATAC-seq measure the genome-wide chromatin accessibility and can locate distal and proximal *cis*-regulatory elements (CREs) (14), and we thus focus on these two technologies in this study.

With the advanced technologies, massive amounts of chromatin accessibility sequencing data has been generated by consortiums or individual research groups and accumulated in repositories such as GEO (15), paving the way to the annotation of chromatin ‘openness’, i.e., the accessibility of genomic regions. Annotating openness of genomic regions of interest is an essential but cumbersome and time-consuming task that involves several manual and tedious steps: (i) identifying the relevant chromatin accessibility experiments, (ii) downloading the large raw sequencing data files, and (iii) calculating the count of reads falling into or peaks overlapping with each genomic region iteratively. Although there are several existing databases storing chromatin accessibility data, such as ATACdb (16), Cistrome DB (17,18), DeepBlue (19), ENCODE (12,13), Ensembl (20), GTRD (21), and OCHROdb (https://dhs.ccm.sickkids.ca/), the comprehensiveness of data collection in most of these databases is insufficient. For example, ATACdb only supports human ATAC-seq data. OCHROdb only supports human DNase-seq data. Both ENCODE and DeepBlue focus on DNase-seq data while containing only a small amount of ATAC-seq samples. Besides, databases providing only open chromatin regions or peaks, such as Cistrome DB, Ensembl and GTRD, hamper the utilization of raw sequencing reads for chromatin accessibility annotation. In addition, existing databases have mainly focused on data collection, whereas efforts to systematically utilize the chromatin accessibility data for annotating openness in batch for a set of genomic regions are limited. Recent toolkits such as Ensembl, Cistrome Toolkit (18) and SCREEN (22) offer the functionality to query a given genomic region. However, the annotation of chromatin accessibility by these toolkits is hindered by the inconvenient use of results for quantitative analysis and, particularly, the inefficient annotation due to the manner of one region per query.

To circumvent these bottlenecks, we developed OpenAnnotate for efficiently annotating openness of massive genomic regions based on comprehensive chromatin accessibility data and facilitating the understanding of regulatory mechanisms (Figure 1). We collected sequencing samples from ENCODE (12,13,23) and ATACdb (16), and categorized them into tissues and biological systems hierarchically. The openness scores are rigorously defined by (i) raw read openness: the normalized number of reads that fall into the regions, and (ii) peak openness: the normalized number of peaks that overlap with the regions. With ultra-efficient annotation and user-friendly functionalities, OpenAnnotate has been successfully applied in revealing cell type-specificity (24), scoring cell type-specific impacts of noncoding variants (25), identifying regulatory elements (26) and 3D chromatin contacts (27), deciphering gene functional relationships by gene co-opening networks (28), inferring functions of transcription factors (29), and unprecedentedly promoting the analyses of single-cell data (30). By detailed tutorials and demos of various applications, we expect that OpenAnnotate will provide a promising avenue for researchers to construct a more holistic perspective to understand regulatory mechanisms.

**Figure 1.**
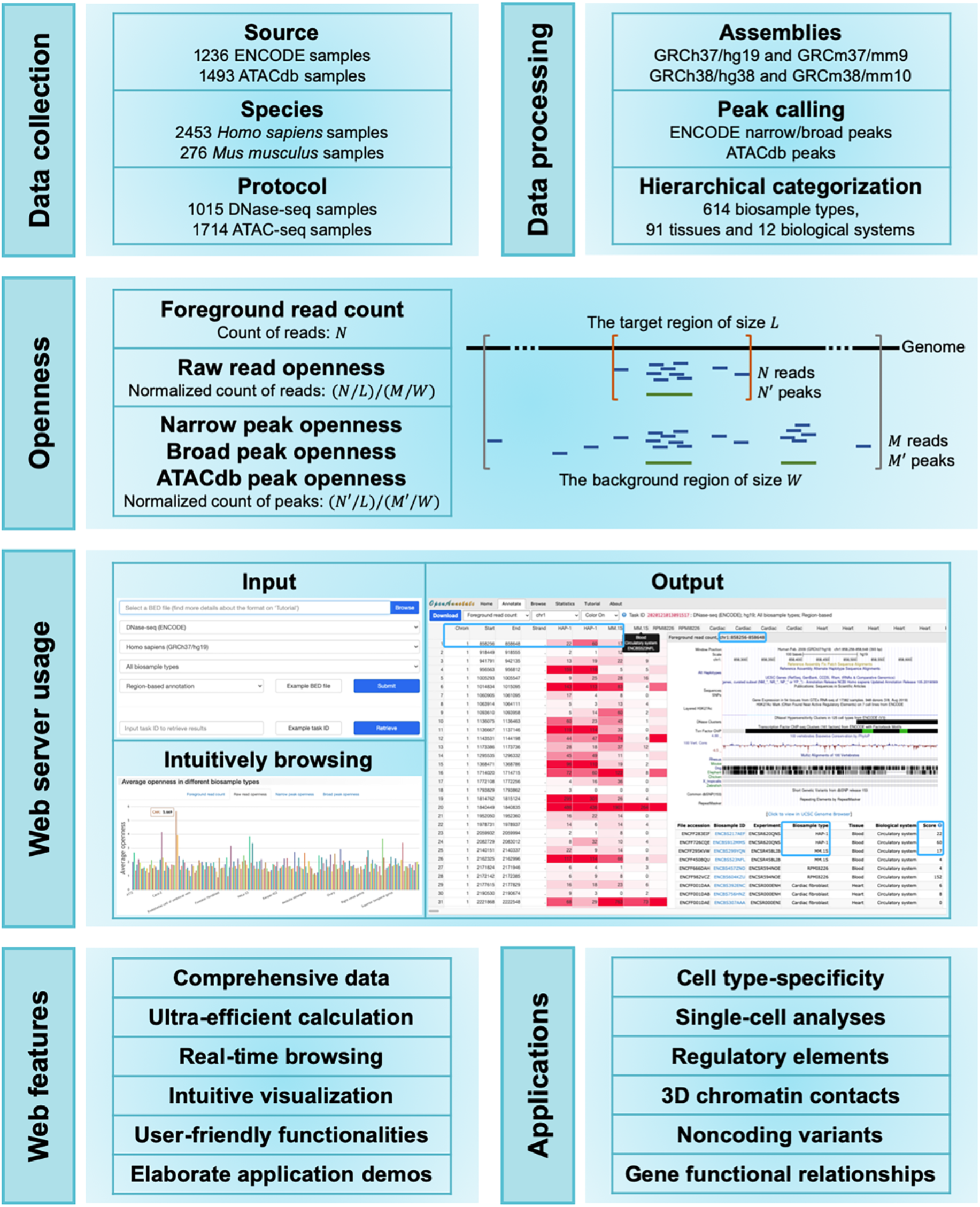
Schematic representation of the OpenAnnotate workflow. Web features and application scenarios are depicted at the bottom.

## MATERIAL AND METHODS

### Data collection and processing

For the openness annotation of genomic regions, we collected a total of 2729 chromatin accessibility samples. As shown in Table 1, 1236 samples of DNase-seq (82.1%) and ATAC-seq (17.9%) were downloaded from ENCODE, a large collaborative project that mainly focuses on providing comprehensive DNase-seq data rather than other chromatin accessibility sequencing data (12,13,23), and 1493 ATAC-seq samples were obtained from ATACdb, the latest chromatin accessibility database which aims to collect a large number of human ATAC-seq data (16). All collected samples were mapped to a specific reference genome (i.e., GRCh37/hg19 for *Homo sapiens* and GRCm37/mm9 for *Mus musculus*). In addition to BAM/SAM files, we also collected peaks that are enriched with aligned reads: (i) for ENCODE data, narrow/broad peaks (peaks/regions of signal enrichment based on pooled and normalized data) identified by the ENCODE analysis pipeline, and (ii) for ATACdb data, peaks identified by the uniform processing pipeline in ATACdb. We then converted the genome coordinates of BAM and BED files from GRCh37/hg19 and GRCm37/mm9 to GRCh38/hg38 and GRCm38/mm10, respectively, using CrossMap (31) with UCSC chain files (32). To facilitate the data traceability, we also provided the original biosample ID in the source of each sample (e.g., ENCBS217AEF in ENCODE and Sample_0001 in ATACdb).

**Table 1.** Statistics of chromatin accessibility data in OpenAnnotate.

| Species | Protocol | Source | Number of samples | Number of biosample types | Number of tissues | Number of biological systems |
| --- | --- | --- | --- | --- | --- | --- |
| *Homo sapiens* | DNase-seq | ENCODE | 871 | 199 | 47 | 11 |
|  | ATAC-seq | ENCODE | 89 | 29 | 24 | 9 |
|  | ATAC-seq | ATACdb | 1493 | 310 | 44 | 12 |
| *Mus musculus* | DNase-seq | ENCODE | 144 | 37 | 18 | 10 |
|  | ATAC-seq | ENCODE | 132 | 66 | 12 | 7 |

We further manually annotated each chromatin accessibility sample with extensive details for hierarchical categorization. For ENCODE samples, a list of biosample types was first extracted from the source information provided by ENCODE, and then manually classified into different tissues. For ATACdb samples, biosample types and tissues were directedly obtained from the metadata annotated by ATACdb. In total, the chromatin accessibility samples were derived from 614 biosample types and 91 tissues, and were finally grouped into stem cell and 10 different biological systems. Samples without enough source information may be ubiquitous and were recorded as ‘Multiple systems’. The detailed statistics of samples in different species, including the number of samples, biosample types, tissues, and biological systems from different protocols and sources, are shown in Table 1.

### Openness annotation

To annotate the openness of each genomic region in each biological sample, OpenAnnotate first calculates the foreground read count (*N*), that is, the number of reads falling into the region using BAM/SAM file of the sample. To correct the effect of sequencing depth, the raw read openness (*S*) is defined as the average foreground read count in the region divided by the average number of reads falling into a background region of size *W* surrounding the given region, and can be calculated as:

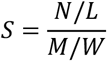

where *L* denotes the length of the given genomic region, and *M* the number of reads falling into the background region. The background-size *W* is set to 1 million base pairs.

Analogously, OpenAnnotate annotates narrow/broad peak openness for ENCODE samples and ATACdb peak openness for ATACdb samples with the average number of peaks overlapping with the given region of size *L* divided by the average number of peaks overlapping with a background region of size *W* surrounding the given genomic region. Together with the two openness scores based on reads and peaks respectively, OpenAnnotate also provides the foreground read count (*N*) for alternative usage. More discussions on the choice of different openness annotation approaches are provided in the APPLICATION SCENARIOS section.

### Implementation

We designed a parallel strategy and deployed OpenAnnotate on a high-performance computing cluster to significantly decrease the time for openness annotation of massive genomic regions in a vast amount of chromatin accessibility samples (Supplementary Note S1 and Supplementary Figure S1). Considering the annotation results can be large tables and loading the entire tables directly to the web is problematic, we adopted WebSocket (https://www.websocket.org/), which provides full-duplex communication channels between web and host, to enable real-time browsing of any part of the resulting large tables. PHP v7.4.5 (http://www.php.net/) was used for server-side scripting, while Bootstrap v3.3.7 framework (https://getbootstrap.com/docs/3.3/) was adopted for the optimization of web-frontend interface. jQuery Plugins and JavaScript Libraries, including DataTables v1.10.19 (https://datatables.net), CanvasJS v2.3.1 (https://canvasjs.com/), and morris.js v0.5.0 (https://morrisjs.github.io/morris.js/index.html), were used to implement advanced tables and responsive charts. The current version of OpenAnnotate runs on a Linux-based Apache web server v2.4.6 (https://www.apache.org) and supports most mainstream web browsers, such as Google Chrome, Firefox, Opera, Microsoft Edge, and Apple Safari.

## WEB SERVER USAGE

OpenAnnotate provides typical applications and elaborate notebooks on the *Home* page, ultra-efficient openness annotation for massive genomic regions on the *Annotate* page, intuitive understanding of the openness of a particular genomic region across various biosample types on the *Browse* page, detailed information and statistics of all the samples on the *Statistics* page, extensive application scenarios and step-by-step tutorials on the *Tutorial* page, and high-lighted information and features of the web server on the *About* page.

### Input

The major functionality of OpenAnnotate is maintained on the *Annotate* page, which requires a tab-delimited BED file containing genomic regions to be annotated (with 1 GB maximum file size restriction). Each line in the BED file represents a genomic region, and has four required fields: the first three fields and the sixth field, defining the name of chromosome, the starting position, the ending position and the strand, respectively. To ensure the number of fields per line is consistent, the strand and other additional column information can be filled with a if the field content is unknown or empty. The BED file can be further compressed in gzip format to accelerate the uploading. Exemplary input files are available for facilitating the use of OpenAnnotate. We also provide users with flexible options to specify the protocol, source, species, assembly, and biosample type for openness annotation. Details about the optional biosample types can be accessed on the *Statistics* page, which provides (i) the metadata of chromatin accessibility samples in a table with advanced features, including searching, sorting, paging, copying, and exporting, and (ii) intuitive comparison of the number of samples across different biosample types, tissues, and biological systems. Furthermore, users can choose the *Per-base pair annotation* mode to calculate the openness of each base-pair of genomic regions for bioinformatics analyses such as machine learning tasks demonstrated in the APPLICATION SCENARIOS section.

Users can obtain a task ID immediately after submitting an annotation task. This unique identifier can be used to retrieve annotation results. We also provide an exemplary task ID to illustrate the usage. Users can also optionally and conveniently archive the task information to a mailbox, including links for checking annotation status and downloading annotation results.

### Output

After submitting the annotation task, users can directly browse real-time annotation results within about 10 seconds, depending on the input file size and internet speed. Corresponding to different annotation approaches, the annotation results are represented as tables where each row represents a genomic region and each column represents the biosample type of a sequencing sample (details can be obtained by hovering over the column name). For browsing, the submitted genomic regions will be grouped into different chromosomes, and the annotation status in different chromosomes will be updated in real-time. Users can arbitrarily switch between different chromosomes or different annotation approaches to browse resulting tables. Although the tables may contain thousands of columns and billions of rows especially on the *Per-base pair annotation* mode, users can browse any part of a large table by scrolling to any row and any column smoothly. By enabling the *Color* option, each element in the table will be colored according to the openness score for intuitively comparing the openness of different genomic regions in various sequencing samples.

Because biosample types are used as column names to provide an intuitive understanding of biosample type-specificity of open regions, there exist duplicate column names for the biosample types with multiple replicates. More detailed information about the sequencing samples (columns), including original biosample IDs in source, categorized biosample types, tissues and biological systems, can be obtained by double-clicking on a row in the table. For the genomic region of the double-clicked row, we also provide the option for visualization in UCSC Genome Browser (32) and the openness scores in various sequencing samples. After the annotation task is completed, users can download the standardized results via the *Download* button, and verify the integrity of a downloaded file with the corresponding MD5 checksum code. Note that the annotation results for real-time browsing are grouped by chromosomes while the annotation results for downloading are maintained in the original order in uploaded BED file.

### Intuitively browsing

To study the openness of a particular genomic region more intuitively, the *Browse* page provides uniform comparisons of openness across the hierarchically categorized samples. After submitting the specified protocol, source, species, assembly, and genomic region, OpenAnnotate offers (i) average openness scores in different biological systems, tissues, and biosample types for intuitive comparison, and (ii) default visualization of the genomic region together with a hyperlink to customize more annotation tracks in UCSC Genome Browser (32).

## APPLICATION SCENARIOS

### OpenAnnotate reveals cell type-specificity of regulatory elements

The openness annotated by OpenAnnotate provides valuable cell type-specific patterns and has been successfully applied to demonstrate the cell type-specificity of validated silencers (24) and model the openness dependence of a regulatory element on its underlying DNA sequence and TF expression for scoring the cell type-specific impacts of noncoding variants in personal genomes (25). Following the analysis pipeline in SilencerDB (24), we demonstrate that OpenAnnotate can also reveal the cell type-specificity of human A549 enhancers collected from EnhancerAtlas 2.0 (33) (Supplementary Note S2).

### OpenAnnotate facilitates studies of regulatory mechanism

OpenAnnotate has been successfully applied to identifying regulatory elements (26) and 3D chromatin contacts (27), deciphering gene functional relationships (28), and inferring functions of transcription factors (29).

#### Identification of regulatory elements

A deep convolutional neural network named DeepCAPE has been proposed to accurately predict enhancers via the integration of DNA sequences and DNase-seq data (26), with the understanding that DNase I hypersensitivity is important to identify active cis-regulatory elements including enhancers, promoters, silencers, insulators, and locus control regions (34). DeepCAPE consistently outperformed other methods, especially those use DNA sequences alone (35), for the identification of cell line-specific enhancers (Supplementary Note S3). Such machine learning frameworks can also be adapted for the prediction of other functional elements to establish a landscape of functional elements specific to a biosample type that still lacks enough systematic exploration.

#### Identification of 3D chromatin contacts

A bootstrapping deep learning model named DeepTACT has been proposed to integrate DNA sequences and chromatin accessibility data for the prediction of chromatin contacts between regulatory elements (27), indicating that the openness annotated by OpenAnnotate plays a crucial role in understanding regulatory mechanisms (Supplementary Note S3).

#### Co-opening analysis

The genes related to a specific biological process tend to be clustered together in gene-gene co-opening networks, which facilitates the elucidation of gene functional relationships (28). Besides, co-opening associations between regulatory regions and nearby genes can provide an accurate interpretation of transcription factor functions, paving the way to elucidate functions of regulatory elements (29).

### OpenAnnotate sheds light on single-cell data analyses

Bulk chromatin accessibility sequencing data has been successfully used as reference data to facilitate characterization (30,36) and imputation (37) of single-cell chromatin accessibility sequencing (scCAS) data. Here, we introduce a straightforward but practical approach, named refProj, to incorporate the openness annotated by OpenAnnotate as a reference for characterizing scCAS data (Supplementary Note S4). Using the source code obtained from a recent benchmark study (38,39), we evaluated the effectiveness of refProj on a scCAS dataset of human hematopoietic cells (referred as Buenrostro2018, Figure 2A) (36). As shown in Figure 2B, refProj-peak (peak openness-based) and refProj-read (read openness-based) achieve the third-best and the sixth-best clustering performance, respectively. Among the top-six methods, SnapATAC (40), Cusanovich2018 (41–43) and cisTopic (44) were shown to be state-of-the-art methods in the benchmark study (38). Data visualization using the top-six methods further demonstrates the effectiveness of refProj to characterize cell heterogeneity (Figure 2C). The results suggest that the openness annotated by OpenAnnotate can facilitate scCAS data analyses (Supplementary Note S4).

**Figure 2.**
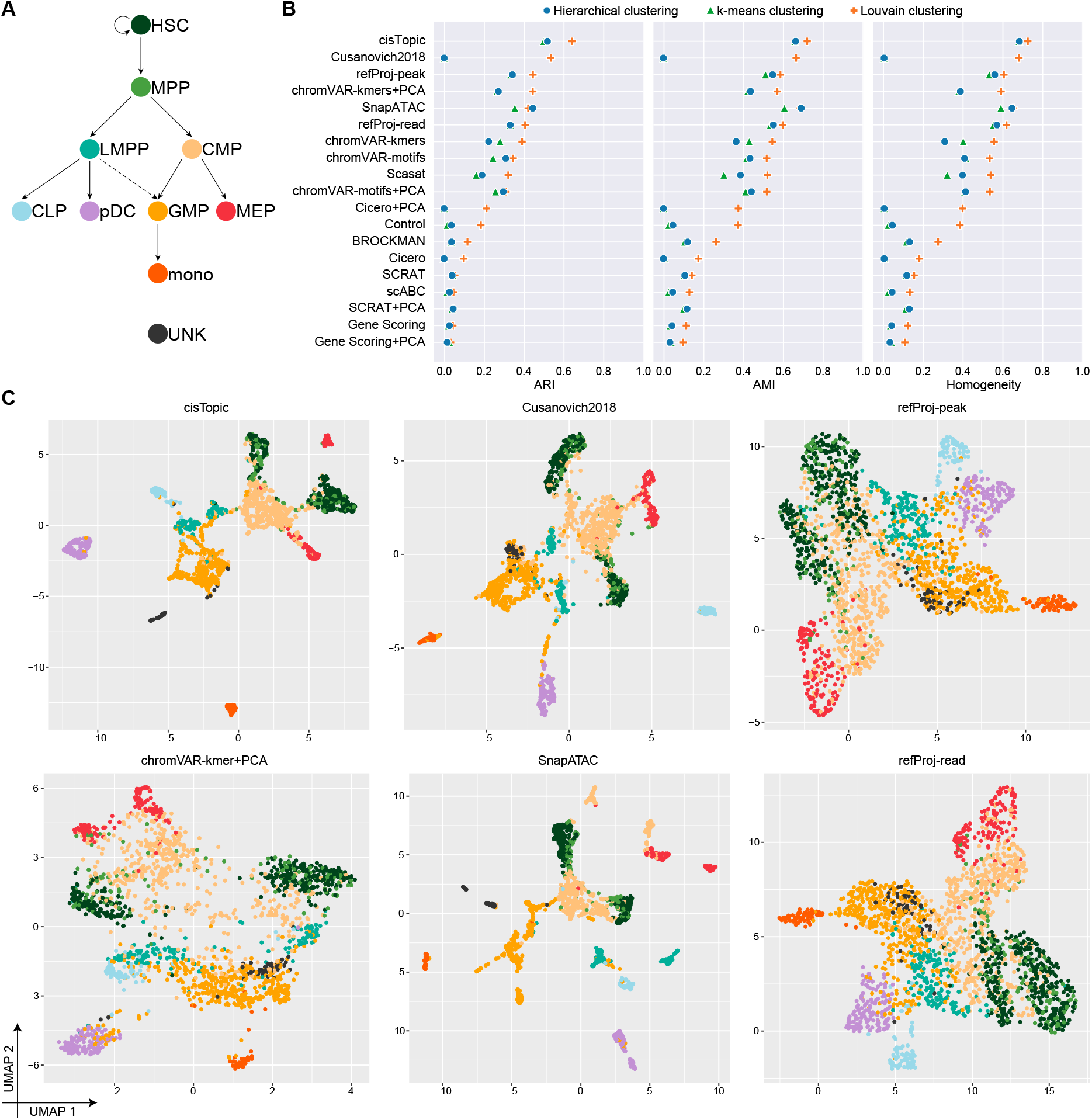
OpenAnnotate facilitates the analysis of scCAS data. (A) Developmental roadmap of cell types in the Buenrostro2018 dataset. (B) Dot plot of scores for each metric to quantitatively measure the clustering performance of each method, sorted by maximum adjusted Rand index (ARI) score. (C) UMAP visualization of the feature matrix produced by the top-six methods ranked in (B). Individual cells are colored indicating the cell type labels shown in (A).

### Selection of the openness annotation approaches

The above examples demonstrate the extensive application scenarios of OpenAnnotate, which aims to efficiently annotate raw read openness, peak openness and foreground read count for massive genomic regions. We note that raw read openness can provide holistic information of chromatin accessibility samples. In contrast, peak openness usually has lower noise but higher sparsity than raw read openness because it is based on peak regions enriched with aligned reads. Raw read openness is the most commonly used annotation, especially on the *Per-base pair annotation* mode because machine learning methods such as convolutional neural networks have an appetite for dense data. To facilitate the analysis of scCAS data in a reference-based manner, peak openness can provide comparable or even better performance than raw read openness, especially when there are massive peaks in the scCAS data (30). Foreground read count can be used in certain cases, such as customizing the normalization method. Besides, cisTopic has presented advantages for the analysis of scCAS data (38,44), indicating that reference-based topic models that directly model foreground read count data can be developed to further promote single-cell data analyses.

## CONCLUSIONS AND FURTHER DIRECTIONS

Chromatin accessibility profiles accumulated in public repositories serve as a valuable resource for studies of regulatory mechanisms. Recent research has mainly focused on data collection, whereas efforts to systematically utilize the data to annotate accessibility of genomic regions are limited. OpenAnnotate is a web server for efficient chromatin accessibility annotation in batch of genomic regions. With comprehensive data collection of 2729 sequencing samples, hierarchical categorization, user-friendly functionalities, ultra-efficient calculation, real-time browsing, intuitive visualization, extensive successful applications, detailed tutorial, as well as step-by-step demos, OpenAnnotate shows numerous advantages compared to existing databases and toolkits (Supplementary Table S1).

OpenAnnotate has five main application scenarios. First, biologists without sophisticated programming skills can conveniently and intuitively compare the openness of genomic regions or a specific region across various biosample types, thus facilitating the study of functional implications (24). Second, computer scientists are currently keen on applying advanced models to solve biological problems. They can use OpenAnnotate to circumvent cumbersome data collection and processing, and incorporate openness data into computational models to introduce cell type-specificity and achieve superior performance (26,27). Third, high degree of sparsity and high technical variation constitute the major hindrances to characterize single-cell chromatin accessibility sequencing (scCAS) data. The bulk openness of peaks in scCAS data can be incorporated as a valuable reference to facilitate the analysis of scCAS data (30,36,37). Fourth, OpenAnnotate offers the potential to reinterpret abundant genetic data from large-scale genome-wide association studies (GWAS). Users can interpret the functional implication of variants by integrating upstream openness and downstream gene expression (25). Finally, the openness annotated by OpenAnnotate can be used for constructing co-opening networks and associations, which provide a new perspective to system biology studies (28,29).

In future versions, OpenAnnotate will follow two main directions. First, we will continue to improve the interfaces and functionalities of OpenAnnotate. Second, we can aggregate cells of the same cell type as a pseudo-bulk sample and incorporate them into our web server. This will make full use of the exponentially accumulated scCAS data and further extend the scope of OpenAnnotate. We anticipate that OpenAnnotate will benefit both biologists and data scientists to better model the regulatory landscape of genome.

## DATA AVAILABILITY

The web server is freely available at http://health.tsinghua.edu.cn/openannotate/ or http://bioinfo.au.tsinghua.edu.cn/openannotate/. This website is open to all users and there is no login requirement. Source code and tutorials for compiling and executing the OpenAnnotate program are freely available at https://github.com/RJiangLab/OpenAnnotate.

## ACKNOWLEDGEMENT

The authors thank Wanwen Zeng and Wenran Li for helpful suggestions.

